# Securin and cyclin B1-CDK1, but not SGO2, regulate separase activity during meiosis I in mouse oocytes

**DOI:** 10.1101/2025.01.06.631516

**Authors:** Benjamin Wetherall, David Bulmer, Alexandra Sarginson, Christopher Thomas, Suzanne Madgwick

## Abstract

During meiosis I in oocytes, anaphase is triggered by deactivation of cyclin B1-CDK1 and activation of separase. Active separase plays an essential role in cleaving cohesin rings that hold homologous chromosomes together. Critically, separase must be inhibited until all chromosomes are aligned and the cell is prepared for anaphase I. Inhibition can be mediated through the binding of separase to either securin or cyclin B1-CDK1. The relative contribution of each inhibitory pathway varies depending on cell type. Recently, shugoshin-2 (SGO2) has also been shown to inhibit separase in mitotic cells. Here, we used a separase biosensor and perturbed the three inhibitory pathways during meiosis I in mouse oocytes. We show that inhibition mediated by either securin or cyclin B1-CDK1, but not SGO2, is independently sufficient to suppress separase activity. However, when both the securin and cyclin B1-CDK1 inhibitory pathways are perturbed together, separase activity begins prematurely, resulting in gross segregation defects. Furthermore, we characterised shugoshin-2 destruction dynamics and conclude that it is not an essential separase inhibitor in mouse oocytes. The existence of multiple separase inhibitory pathways highlights the critical importance of tightly regulated separase activity during this unique and challenging cell division.

## Introduction

Meiosis is the specialised cell division that generates female and male gametes, the egg and sperm. In meiosis I, chromosomes are organised in a structure known as a bivalent. The bivalent is composed of two homologous chromosomes of different parental origin^1^. Both homologous chromosomes within the bivalent, as well as the sister chromatids within each homolog, are held together by a ring-like protein complex known as cohesin^2^.

For correct progression through the meiotic divisions, cohesin must be removed in distinct steps. In meiosis I, cohesin is cleaved from chromosome arms - allowing the homologous chromosomes within each bivalent to segregate. In meiosis II, cohesin still present at the pericentromeric regions of chromosomes is cleaved - allowing sister chromatids to separate^3^. In both divisions, the protease separase plays an essential role, by cleaving the meiosis-specific kleisin subunit of cohesin – REC8^4–6^. It is therefore critical that separase is carefully regulated through both divisions and only becomes active once chromosomes are aligned and prepared to divide.

In mouse oocytes, two inhibitory pathways have been implicated in separase inhibition. In the first pathway, separase directly interacts with securin, which acts as a pseudosubstrate inhibitor of separase^7–13^. Additionally, upon phosphorylation by CDK1, separase is inhibited by binding CDK1’s activating partner cyclin B1^14–17^. The binding of separase with either securin or cyclin B1 is mutually exclusive, and the relative contribution of each pathway varies depending on cell type and developmental state^15,17^. For example, in female mouse meiosis II and eukaryotic mitosis, securin is largely responsible for separase inhibition, while primordial germ cells and early-stage embryos rely primarily on cyclin B1-CDK1-mediated separase inhibition^18–20^. Interestingly however, while securin binding to separase is the primary inhibitory mechanism in mitotic cells, it is also dispensable, demonstrating the compensatory nature of these two pathways^21–23^. Similarly, in meiosis I mouse oocytes, Chiang et al. only observed chromosome segregation errors when both securin- and cyclin B1-CDK1-mediated separase inhibition were perturbed together^24^. However, in this study, separase activity was not measured directly, and it remains unclear when and how these segregation defects arise.

Shugoshin-2 (SGO2) has also recently been shown to have the capacity to inhibit separase in mitotic cells^25^. This work demonstrates that on association with spindle assembly checkpoint (SAC)-activated MAD2, SGO2 can sequester the majority of free separase in securin knock-out cells. Like securin, through a pseudosubstrate sequence within its N-terminus, SGO2 can bind directly to the active site of separase, blocking its protease activity. It is currently unknown whether SGO2-MAD2 contributes to separase inhibition during oocyte meiosis. When we consider that securin and cyclin B1-CDK1 are regulated differently in oocytes compared to mitosis^26,27^, it is important to investigate whether SGO2 might contribute to separase inhibition during this specialised division.

Therefore, while we have some understanding of the pathways regulating separase in mouse oocytes, exactly how these proteins work together to keep separase activity at bay through the extended duration of meiosis I remains unclear.

In this study, we set out to address the question of separase regulation in mouse oocytes using a live-cell imaging approach. We utilised a fluorescent biosensor to visualise separase activity, in combination with strategies to perturb key cell cycle proteins of interest. We add dynamic understanding to previous findings that either securin- or cyclin B1-CDK1-mediated inhibition are independently sufficient to suppress separase activity in oocyte meiosis I. When both compensatory pathways are disrupted in tandem, separase activity initiates up to 1 hour ahead of anaphase I and is accompanied by gross segregation defects. We additionally investigated the destruction dynamics of SGO2 in mouse oocytes and determined that it is unlikely to function as a separase inhibitor in unperturbed oocytes. The existence of multiple layers of redundancy and crosstalk between separase inhibitory pathways emphasises the crucial role of precisely regulated separase activity in coordinating oocyte meiosis successfully.

## Results

### Both securin- and cyclin B1-CDK1-mediated inhibition are independently sufficient to suppress separase activity in meiosis I

We set out to investigate the pathways regulating separase activity in mouse oocytes. To do this, we used strategies to inhibit the known meiotic separase inhibitors securin and cyclin B1-CDK1 to assess their contributions. Chiang et al. previously showed that segregation errors in fixed meiosis II oocytes were only observed when both securin- and cyclin B1-CDK1-mediated separase inhibition were removed^24^. However, when these errors arise and how separase is regulated through time remain unclear. To investigate this, we used a live separase activity biosensor generated by Nam et al. which has been validated in HeLa cells, U2OS cells, mouse embryonic fibroblasts (MEFs), and mouse oocytes^28–31^. The sensor is composed of a nucleosome-targeted H2B protein that is fused to eGFP and mCherry fluorophores. Active separase cleaves an Scc1 peptide sequence located between the two fluorophores (Fig. 1a). This cleavage leads to a shift in fluorescence from yellow to red, as the eGFP signal dissociates into the cytoplasm while mCherry remains attached to histones linked to chromosomal DNA (Fig. 1b). We chose to use the Scc1 separase biosensor due to the pronounced fluorescence change, making quantification of cleavage timings most reliable. However, we first confirmed that the Scc1 sensor cleavage takes place with the same timing as the Rec8 sensor in mouse oocytes (Fig. 1b).

**Figure 1.**
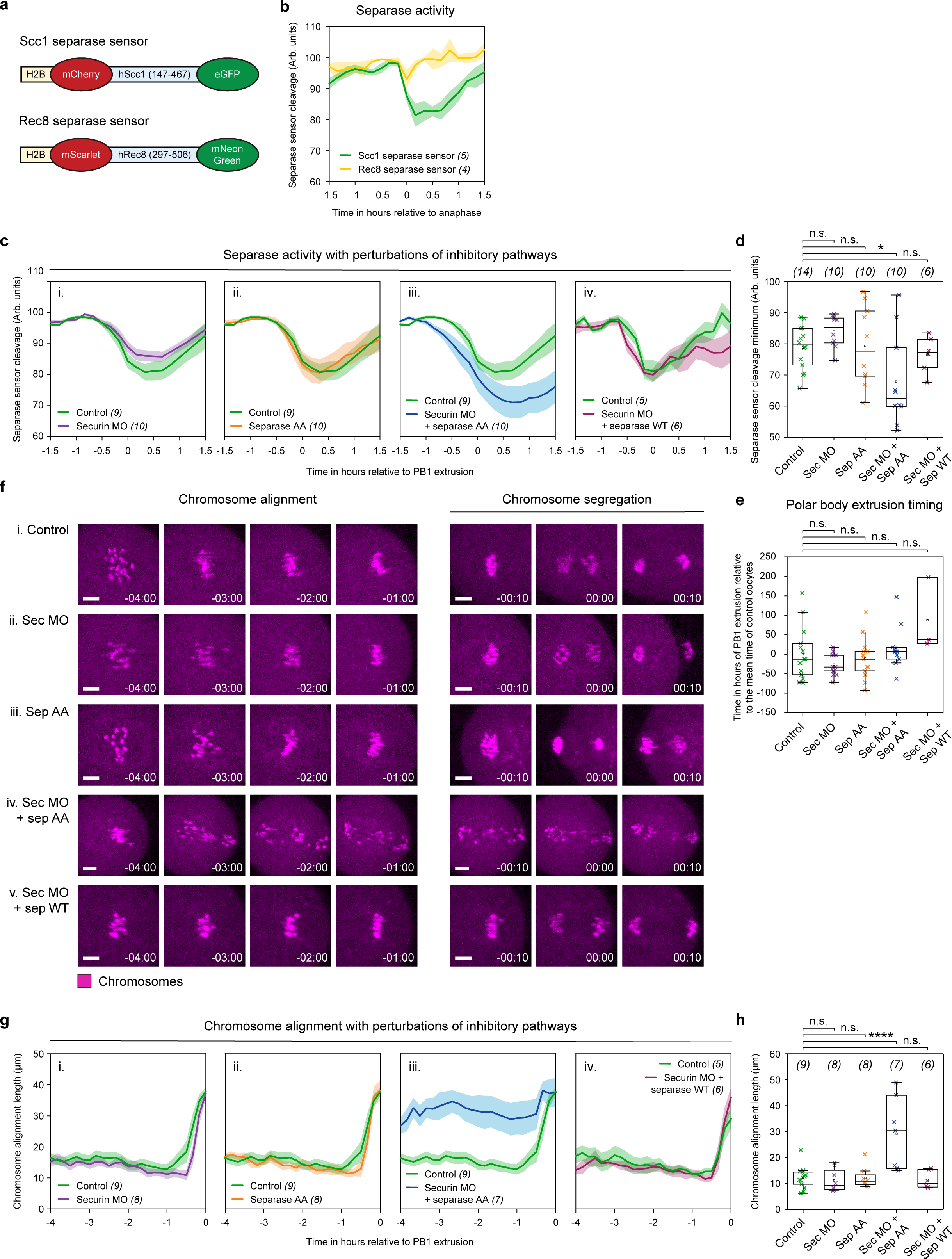
Both securin- and cyclin B1-CDK1-mediated inhibition are independently sufficient to suppress separase activity in meiosis I. **(a)** Schematic diagrams of H2B-mCherry-hScc1-eGFP and H2B-mScarlet-hRec8-mNeonGreen separase activity biosensors. **(b)** Graph comparing Scc1 (green trace, n = 5) and Rec8 (yellow trace, n = 4) separase sensor detection in oocytes aligned at PB1 extrusion. Graphs showing mean MI separase activity profiles as determined by the Scc1 separase sensor (eGFP/mCherry ratio) for control oocytes (green trace, n = 9) compared to (i) securin MO oocytes (purple trace, n = 10) (ii) separase AA oocytes (orange trace, n = 10) and (iii) securin MO + separase AA oocytes (blue trace, n = 10) . To control for overexpression of separase in (iii), (iv) directly compares securin MO + separase WT oocytes (burgundy trace, n = 6) with control oocytes (green trace, n = 5). Quantification of separase sensor cleavage minimum in control (green crosses, n = 14), securin MO (purple crosses, n = 10), separase AA (orange crosses, n = 10), securin MO + separase AA (blue crosses, n = 10), and securin MO + separase WT (burgundy crosses, n = 6) oocytes. Crosses represent individual oocytes. **(e)** Quantification of polar body extrusion timings in oocyte treatment groups as per part (c-d). Timings are presented relative to the mean polar body extrusion time (0) in control oocytes. **(f)** Representative images showing chromosome alignment and segregation in (i) control, (ii) securin MO, (iii) separase AA, (iv) securin MO + separase AA, and (v) securin MO + separase WT oocytes. Time relative to PB1 extrusion. Chromosomes were visualised by incubating oocytes with SiR-DNA (magenta). Scale bar = 10 µm. **(g)** Graphs showing the maximum distance between chromosomes, measured axial to the spindle per 10-minute time points over 4 hours prior to PB1 extrusion. Comparing control oocytes (green traces, n = 9) with (i) securin MO (purple trace, n = 8), (ii) separase AA (orange trace, n = 8), (iii) securin MO + separase AA (blue trace, n = 7) and (iv) securin MO + separase WT (burgundy trace, n = 6) oocytes. **(h)** Quantification of chromosome alignment length in individual oocytes 1 hour before PB1 extrusion in treatment groups as in panel (g). Crosses represent individual oocyte measurements. In panels: (b), (c) and (g): thick lines show the mean data, lighter shadows show the SEM. In panels (d), (e) and (h): **** = P<0.0001, *** = P<0. 001, ** = P<0.01, * = P<0.1, n.s. = non-significant, two-sided unpaired t-test; box = 25-75%, whiskers = 0-100%, centre = median, centre box = mean.

We first used the Scc1 separase sensor in combination with knockdown of securin by morpholino oligo (MO) such that in prometaphase I, MO oocytes contained ∼13% of the protein level relative to non-treated control oocytes (as previously quantified by western blot^26^). Despite severe depletion of securin, the cleavage activity of separase was still restricted to the final 30 minutes before polar body extrusion (Fig 1ci). This result agrees with observations made in fixed oocytes, suggesting that cyclin B1-CDK1 is sufficient to compensate for the loss of securin^24^. To test whether the reverse is also true, we injected oocytes with a previously published separase AA mutant^24^. This construct is fully active but lacks CDK1 phosphorylation sites (S1121A and T1342A) that prevent its inhibition by cyclin B1-CDK1. By this strategy, cyclin B1-CDK1-mediated separase inhibition is perturbed without affecting other crucial functions of cyclin B1-CDK1 in meiosis. Importantly, oocytes expressing separase AA produce separase activity profiles identical to control oocytes, suggesting that securin was similarly sufficient to compensate for a loss of cyclin B1-CDK1-mediated inhibition (Fig. 1cii). In contrast, where securin protein levels are restricted in separase AA expressing oocytes (securin MO + separase AA), cleavage activity is detected up to 1 hour ahead of control oocytes (Fig. 1ciii). Furthermore, securin MO + separase AA oocytes have a significantly greater total cleavage compared to control oocytes (Fig. 1d).

The difference in cleavage timing is also apparent in the confocal microscopy images, where the eGFP signal remains high on chromosomes in control oocytes until just prior to chromosome segregation. In contrast, in securin MO + separase AA oocytes, the eGFP signal begins to visibly decrease approximately 30 minutes earlier (Supplementary Fig. 1b). Importantly, this premature separase activity was specific to the loss of both securin- and cyclin B1-CDK1-mediated inhibition, and not due to separase overexpression in the absence of securin. Oocytes treated with securin MO + an excess of wild-type separase have cleavage timings identical to control oocytes (Fig. 1civ), suggesting that cyclin B1-CDK1 alone has the capacity to inhibit an abundance of separase. Despite the premature separase activity in securin MO + separase AA oocytes, we observed no change in the timing of polar body extrusion (Fig. 1e).

To monitor the impact of premature separase activity in securin MO + separase AA oocytes, we used confocal microscopy to image live oocytes stained with SiR-DNA to track individual chromosome movements. This analysis revealed severe defects in chromosome alignment and segregation in securin MO + separase AA oocytes (Fig. 1fiv). Additionally, quantification of the maximum distance between chromosomes in each 10-minute time interval prior to polar body extrusion showed significantly more disorganised metaphase plates during meiosis I chromosome alignment in securin MO + separase AA oocytes (Fig. 1g). Notably, this analysis also revealed that 1 hour prior to PB1 extrusion, even before detectable cleavage of the sensor begins, the average length of the chromosome plate in securin MO + separase AA oocytes was significantly longer (29.2 ± 5.3 µm; Fig. 1h) than in control oocytes (12.3 ± 1.1 µm; Fig. 1h). In contrast, securin MO (11.2 ± 1.5 µm; Fig. 1h), separase AA (12.2 ± 1.5 µm; Fig. 1h), and securin MO + separase WT (11.4 ± 1.4 µm; Fig. 1h) oocytes were the same as controls.

These data further demonstrate that either inhibitory pathway, securin or cyclin B1-CDK1, is sufficient to prevent premature separase activity in mouse oocyte meiosis I. When both pathways are perturbed in the same oocyte, the timings of anaphase I onset and separase release become uncoupled, resulting in severe defects in chromosome alignment and segregation.

### Shugoshin-2 is targeted for degradation by the APC/C in anaphase I

While the defects observed in securin MO + separase AA oocytes were severe, we found it curious that even when both inhibitory pathways were perturbed, detectable cleavage of the sensor was only observed in the final 1.5 hours before polar body extrusion (Fig. 1ciii). We therefore wanted to investigate whether another inhibitory mechanism may be functioning in these oocytes. Specifically, SGO2, recently shown to be capable of separase inhibition in human mitotic cells when securin was knocked-out^25^.

SGO2 contains a conserved pseudosubstrate sequence within its N-terminus (Fig. 2a) that can directly bind the active site of separase, blocking its protease activity^25^. We hypothesised that SGO2 may be contributing to separase inhibition in securin MO + separase AA oocytes, and the release of separase activity around 1.5 hours ahead of polar body extrusion in these oocytes (Fig. 1ciii) may reflect a time when the inhibitory capacity of SGO2 is lost. We further hypothesised that this loss may be due to SGO2 being targeted for proteasomal degradation by the APC/C, like securin, cyclin B1, and indeed its homolog SGO1^32–34^. To support this hypothesis, when we compared multiple sequence alignments of SGO2, we identified several conserved APC/C destruction motifs^35^ (Fig. 2a). Specifically, the N-terminus of SGO2 contains a previously undescribed KEN box^36^, TEK box^37^, and D-box^38^ just upstream of the separase pseudosubstrate motif identified by Hellmuth et al^25^. To further investigate whether SGO2 is targeted for destruction during meiosis I, we measured SGO2 protein levels in oocytes using a live fluorescent SGO2 reporter. We observed that the SGO2 reporter was rapidly degraded late in meiosis I, only 30 minutes ahead of PB1 extrusion and over an hour later than both securin and cyclin B1 destruction is initiated (Fig. 2b). We then confirmed that the timing of SGO2 degradation is unchanged between control and securin MO + separase AA oocytes (Fig. 2c). This late destruction suggests that SGO2 is not mediating separase inhibition in securin MO + separase AA oocytes, as SGO2 is still present at high levels for over an hour after the point of premature separase activation (Fig. 1ciii).

**Figure 2.**
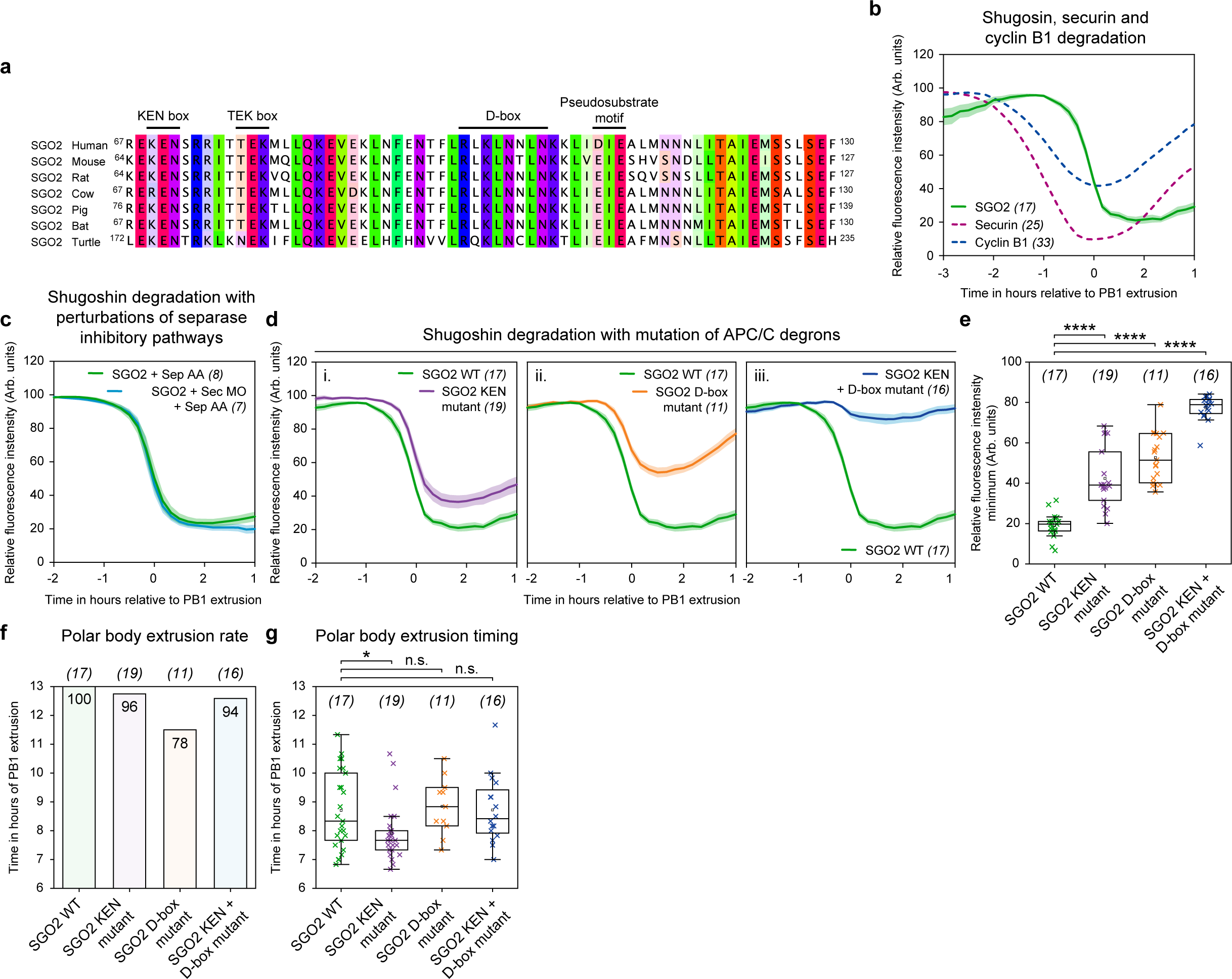
Shugoshin-2 is targeted for degradation by the APC/C in anaphase I. **(a)** Multiple sequence alignment of the N-terminal segments in shugoshin-2 orthologs containing the KEN box, TEK box, D-box, and separase pseudosubstrate motif. **(b)** Mean SGO2 (green trace, n = 17), securin (burgundy dashed trace, n = 25), and cyclin B1 (blue dashed trace, n = 33) destruction profiles relative to PB1 extrusion. **(c)** Mean SGO2 destruction traces comparing oocytes with separase AA expression (green trace, n = 8) and securin MO + separase AA expression (blue trace n = 7). **(d)** SGO WT destruction traces relative to PB1 extrusion (green trace, n = 17) compared to (i) SGO2 KEN mutant (purple trace n = 19), (ii) SGO2 D-box mutant (orange trace, n = 11) and (iii) SGO2 KEN + D-box mutant (blue trace, n = 16). **(e)** Quantification of relative degradation by fluorescence intensity minimum for groups in panel (d). Crosses represent individual oocyte measurements. Plots **(f)** and **(g)** display polar body extrusion rates and polar body extrusion timings respectively for groups in panels (d) and (e). In panels (b), (c) and (-d): thick lines show the mean data, lighter shadows show the SEM. In panels (e) and (g): **** = P<0.0001, *** = P<0. 001, ** = P<0.01, * = P<0.1, n.s. = non-significant, two-sided unpaired t-test; box = 25-75%, whiskers = 0-100%, centre = median, centre box = mean.

Interestingly, when we mutated the KEN box and D-box, which had an additive effect in stabilising SGO2 during anaphase I (Fig. 2d-e), this had no impact on the polar body extrusion rate or timing (Fig. 2f-g). This suggests that unlike in securin and cyclin B1^39^, KEN/D-box mutants of SGO2 are unable to block anaphase onset and polar body extrusion through persistent separase inhibition in mouse oocytes. However, due to the close vicinity of the KEN box and D-box to the separase-interacting pseudosubstrate motif in SGO2 (Fig. 2a), we cannot discount that these mutants fail to block progression into meiosis II due to an impaired capacity to interact with separase. Furthermore, in mitotic cells, release of separase from SGO2-mediated inhibition occurs in an APC/C-independent manner^25^. We therefore looked for an additional approach to assess the impact of SGO2 on separase inhibition in oocytes.

### Shugoshin-2 is not essential for separase inhibition in meiosis I

Using a morpholino oligo (MO), we knocked down SGO2 such that 3 hours after NEBD, MO oocytes contained <15% of the protein level of non-treated control oocytes (as quantified by immunofluorescence; Supplementary Fig. 2a-b). Even with this depletion of SGO2 protein, like in control oocytes, the cleavage activity of separase remained restricted to the final 30 minutes preceding PB1 extrusion (Fig 3ai). We additionally injected oocytes with SGO2 MO + securin MO (Fig. 3aii) and SGO2 MO + separase AA (Fig. 3aiii). In both of these double perturbations, the cleavage timing was consistent with that observed in control oocytes. Together, these data provide important evidence that in unperturbed mouse oocytes, SGO2 does not play an essential role in the inhibition of separase during meiosis I.

We next wanted to revisit the idea that in securin MO + separase AA oocytes, SGO2 may be able to compensate for the loss of securin- and cyclin B1-CDK1-mediated separase inhibition prior to becoming active in the final 1.5 hours before PB1 extrusion. To address this, we knocked down both SGO2 and securin protein levels in separase AA expressing oocytes (SGO2 MO + securin MO + separase AA). In these oocytes, the loss of SGO2 had no further impact on the timing of cleavage activity, which still initiates up to 1.5 hours ahead of PB1 extrusion, like that of securin MO + separase AA oocytes (Fig. 3aiv). However, there was a marked change in the extent to which the sensor was cleaved in SGO2 MO + securin MO + separase AA oocytes, now significantly reduced (Fig. 3b), and accompanied by a delay in PB1 extrusion (Fig. 3c).

**Figure 3.**
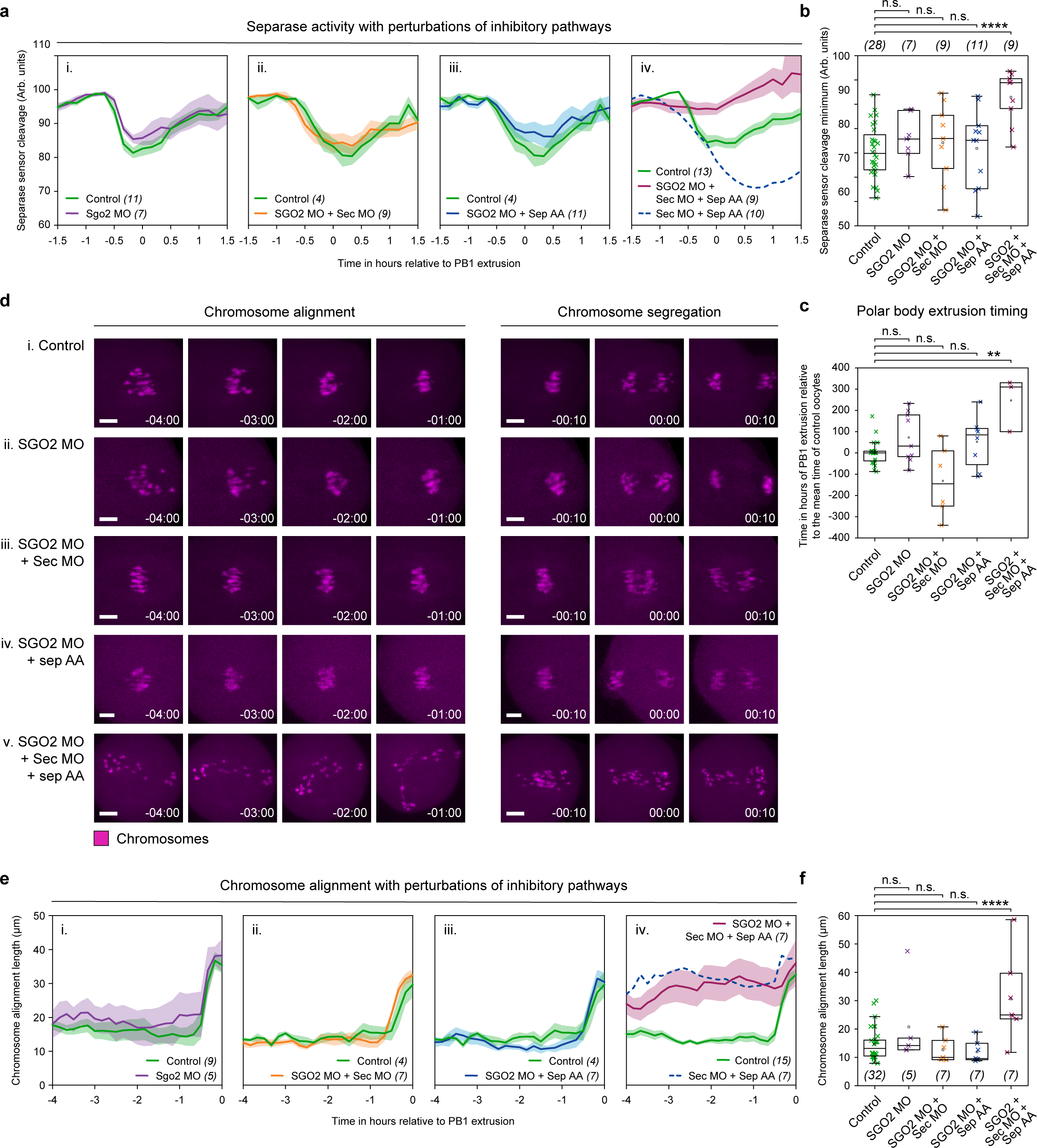
Shugoshin-2 is not essential for separase inhibition in meiosis I. **(a)** Graphs showing mean MI separase activity profiles as determined by the Scc1 separase sensor (eGFP/mCherry ratio) for control oocytes (green trace, n = 11) compared to (i) SGO2 MO (purple trace, n = 7), (ii) SGO2 MO + securin MO (orange trace, n = 9), (iii) SGO2 MO + separase AA (blue trace, n = 11), and (iv) SGO2 MO + securin MO + separase AA oocytes (burgundy trace, n = 9). In part (iv) a dashed dark blue line is included for comparison, indicating the separase sensor ratio in securin MO + separase AA oocytes (n = 10). Quantification of separase sensor cleavage minimum in control (green crosses, n = 28), SGO2 MO (purple crosses, n = 7), SGO2 MO + securin MO (orange crosses, n = 9), SGO2 MO + separase AA (blue crosses, n = 11), and SGO2 + securin MO + separase AA (burgundy crosses, n = 9) oocytes. Crosses represent individual oocytes. **(c)** Quantification of polar body extrusion timings in oocyte treatment groups shown in panel (b). Timings are relative to the mean polar body extrusion time (0) in control oocytes. **(d)** Representative images showing chromosome alignment and segregation in (i) control, (ii) SGO2 MO, (iii) SGO2 + securin MO, (iv) SGO2 MO + separase AA, and (v) SGO2 + securin MO + separase AA oocytes. Time relative to PB1 extrusion. Chromosomes were visualised by incubating oocytes with SiR-DNA (magenta). Scale bar = 10 µm. **(e)** Quantification of chromosome alignment length, measured axial to the spindle per 10-minute time points over 4 hours prior to PB1 extrusion. Comparing control oocytes (green traces, n = 9) with (i) SGO2 MO (purple trace, n = 5), (ii) SGO2 MO + securin MO (orange trace, n = 7), (iii) SGO2 MO + separase AA (blue trace, n = 7) and (iv) SGO2 + securin MO + separase WT oocytes (burgundy trace, n = 7). In part (iv) a dashed dark blue line is included for comparison, indicating chromosome alignment in securin MO + separase AA oocytes (n = 7). **(f)** Quantification of chromosome alignment length 1 hour before PB1 extrusion in treatment groups as shown in panel (e). Crosses represent individual oocyte measurements. In panels (a) and (e): thick lines show the mean data, lighter shadows show the SEM. In panels (b), (c) and (f): **** = P<0.0001, *** = P<0. 001, ** = P<0.01, * = P<0.1, n.s. = non-significant, two-sided unpaired t-test; box = 25-75%, whiskers = 0-100%, centre = median, centre box = mean.

To monitor the impact of premature separase activity in SGO2 MO + securin MO + separase AA oocytes, we imaged live oocytes stained with SiR-DNA to track individual chromosome movements. This analysis revealed errors in chromosome alignment and segregation (Fig. 3dv) comparable to securin MO + separase AA oocytes (Fig. 1fi). In addition, in SGO2 MO + securin MO + separase AA oocytes, the length of the chromosome plate 1 hour prior to PB1 extrusion (30.6 ± 5.6 µm) was double that in control oocytes (14.7 ± 1 µm; Fig. 3e-f) and almost identical to that of securin MO + separase AA oocytes (Fig. 1h).

In conclusion, these findings, along with the destruction timings shown in Fig. 2, provide compelling evidence that SGO2 does not play an essential role in separase inhibition during meiosis I in mouse oocytes.

## Discussion

Accurate progression through meiosis in oocytes requires the timely removal of key cell cycle proteins. From prometaphase I, the removal of cyclin B1 and securin are synchronous, ensuring that CDK1 activity loss is coupled to separase release. Importantly, while the two events are connected, loss of CDK1 activity initiates ahead of separase activation by approximately 30 minutes^27^. This ordering ensures that separase is not released until the cell has committed to anaphase. Aberrant separase release ahead of this time point could have severe consequences. Indeed, in securin MO + separase AA oocytes, separase activity initiates at the same time as the loss in CDK1 activity in unperturbed oocytes, resulting in significant segregation defects^27^.

While cyclin B1-CDK1 drives many diverse processes as a master cell cycle regulator in mitosis and meiosis^40,41^, securin has been shown to primarily function as a separase inhibitor^42^. Both must be depleted to release separase activity during exit from meiosis I^24,39^. Importantly however, cyclin B1 and securin are only partially degraded, and some CDK1 activity remains during meiosis I exit in mouse oocytes^26,27^. This situation is intrinsic to the production of a haploid MII oocyte. Complete cyclin B1-CDK1 activity loss would drive the oocyte into S-phase^43^, while complete loss of securin results in loss of sister chromatid cohesion and subsequent segregation errors^44,45^. Separase must therefore be carefully regulated through the MI-MII transition, perhaps explaining the existence of compensatory mechanisms of separase inhibition.

In the current study we demonstrate that exogenous SGO2 is targeted for degradation by the APC/C in anaphase I, after cyclin B1 and securin, where destruction initiates in prometaphase I. Initially it seemed plausible that SGO2 may act as a late inhibitor of separase in oocytes. However, severe SGO2 depletion alone does not impact the timing of separase release, and SGO2 depletion in securin MO + separase AA oocytes neither alters the timing of sensor cleavage nor exacerbates defects in chromosome alignment and segregation. In addition, it is perhaps counterintuitive for SGO2 to act as a separase inhibitor in mouse oocytes, since direct SGO2-MAD2 binding has previously been demonstrated to be important for SAC silencing during meiosis I^46^. SGO2-MAD2 thereby promotes securin and cyclin B1 destruction to activate, rather than inhibit, separase. A recent study also demonstrated that MAD2 is dispensable for accurate chromosome segregation in mouse oocytes using mice with an oocyte-specific depletion of MAD2^47^. We suggest that in a healthy oocyte, it is not necessary for SGO2 to play a separase inhibitory role, and that in an ageing context, the depletion of SGO2 and MAD2 makes it less likely that any inhibitory contribution would be meaningful^48–50^.

A caveat to our study is that our methods relied on the use of morpholino oligos, which, while severely depleting securin and SGO2, leave some residual protein. Nevertheless, limiting both cyclin B1-CDK1-mediated and securin-mediated separase inhibition accelerates separase release by ∼1 hour. But what inhibits separase before this time point? Notably, initiation of separase release is still acute in this double perturbation context, and this is important. While it seems unlikely that SGO2 takes on this role (SGO2 depletion has no additive impact), it is possible that an unknown separase inhibitor acts over this time period. A characteristic of this inhibitor would be its loss, inactivation or relocalisation ∼1.5 hours ahead of polar body extrusion; currently uncharacterised SGO2 isotypes present possible candidates. However, we propose an alternative and suggest that this is more likely. In securin MO + separase AA oocytes, our data and that of others strongly suggests that endogenous cyclin-CDK1 is sufficient to inhibit all endogenous separase activity. Most convincingly, female mice devoid of securin are fertile^51,52^. In this context, our experimental oocytes are then only vulnerable to exogenous separase AA, which might not be in excess of even very reduced levels of securin in securin MO oocytes. It is plausible that residual securin is sufficient to inhibit separase AA up to a tipping point that is reached earlier in these oocytes when compared to control oocytes with high levels of securin. The period of separase inhibition beyond the initiation of cyclin B1 and securin destruction ∼2.5 hours ahead of polar body extrusion, is then directly governed by their availability. Experiments using securin knockout mice and separase resistant to inhibition by cyclin B1-CDK1 will be well suited to resolve this in future studies.

A second important question raised by our findings is why there are such pronounced alignment defects in securin MO + separase AA (and SGO2 MO + securin MO + separase AA) oocytes, hours before detectable separase activity. In these oocytes, chromosomes are disorganised from initial spindle assembly compared to controls. We suggest that this impact is also related to the addition of separase AA since mRNA is microinjected into GV stage oocytes. At this time point in unperturbed oocytes, endogenous securin and separase levels are low. In our experimental setting, we shift the balance towards substantially reduced securin (due to prior securin MO addition) but generate separase that cannot be inhibited by cyclin B1-CDK1. Following meiosis I resumption, as spindle assembly initiates, it is plausible that there is aberrant early separase activity, which could result in chromosome architecture damage from the outset. This would be difficult to detect, as the H2B-targeted separase sensor does not become discretely localised until after chromosome condensation.

Given its known role in protecting centromeric cohesion in oocytes^53,54^, one might expect that the removal of SGO2 would result in segregation defects. However, this was not detectable in our study. In SGO2-depleted (SGO2 MO) oocytes, as well as in SGO2 MO + securin MO and SGO2 MO + separase AA oocytes, separase activation occurs at the same time as in control oocytes, and chromosomes segregate correctly. Interestingly however, in oocytes where SGO2 was removed in a securin MO + separase AA background (SGO2 MO + securin MO + separase AA), total cleavage of the separase sensor was significantly reduced, and polar body extrusion was delayed. This is in contrast to securin MO + separase AA oocytes, where the total cleavage of the sensor was substantially higher than in control oocytes. We propose that this is due to the previously described additional role of SGO2 in silencing the spindle assembly checkpoint (SAC) during meiosis I in oocytes^55^. In securin MO + separase AA oocytes, the severe errors in chromosome alignment are expected to result in residual high SAC activity later into meiosis compared to controls. When this high SAC activity is coupled with the removal of SGO2, as in SGO2 MO + securin MO + separase AA oocytes, the inability to efficiently deactivate the SAC may hinder APC/C-mediated degradation of securin and cyclin B1. This would result in reduced separase activation, and the reduced sensor cleavage and delayed polar body extrusion observed in these oocytes.

To conclude, our results demonstrate that either securin- or cyclin B1-CDK1-mediated inhibition is independently sufficient to suppress separase during meiosis I in oocytes. In contrast, when these compensatory pathways are perturbed in tandem, separase activity begins up to 1.5 hours ahead of polar body extrusion and is accompanied by gross segregation defects. This erroneous phenotype is not exacerbated when SGO2 protein levels are also depleted. Compensatory pathways of separase inhibition may have multiple advantages, some of which are discussed above but also include changes that take place during oocyte ageing. Here the protein balance in the oocyte shifts towards an increased risk of aberrant separase activity; greater numbers of separase transcripts are observed^56^, while the APC/C becomes more active during anaphase I, destroying greater quantities of securin^44^. In addition, aged oocytes have decreased levels of SAC proteins^44,57–59^, and a diminished population of both SGO2 and cohesin that act to maintain homologous chromosome architecture during meiosis I^60–65^.

This work highlights another example of the oocyte’s ability to adapt to the unique challenges presented during this specialised cell division.

## Acknowledgements

This work was supported by an MRC Career Development Award to S.M. [MR/T010789/1]. We would also like to thank Glyn Nelson (Bioimaging Unit, Faculty of Medical Sciences, Newcastle University) for his invaluable help with imaging.

## Author Contributions Statement

B.W. carried out most experiments alongside C.T. and S.M., with critical contributions from A.S. in immunofluorescence experiments and D.B in confocal microscopy experiments. The project was carried out under the supervision of S.M. C.T. and S.M. designed the experiments, interpreted the data, and prepared the manuscript alongside B.W.

## Supplementary figure legends

## Methods

### Oocyte collection and culture

6-to-12-week-old female, outbred, CD1 mice (Charles River) were used. All animals were handled in accordance with ethics approved by the UK Home Office Animals Scientific Procedures Act 1986. However, given that mice did not undergo a ‘procedure’ as defined by the Act, the project did not require Home Office Licensing. The reason for animal use was instead approved and governed by Newcastle University’s Comparative Biology Centre Ethics Committee; AWERB Approval Reference Number; 663. GV stage oocytes were collected from ovaries punctured with a sterile needle and stripped of their cumulus cells mechanically using a pipette. For bench handling, microinjections and imaging experiments, oocytes were cultured at 37°C in M2 medium (Sigma Aldrich) supplemented with Penicillin-Streptomycin and where necessary with the addition of 30 nM 3-isobutyl-1-methylxanthine (Sigma Aldrich) to arrest oocytes at prophase I. Data were only collected from oocytes that underwent NEBD with normal timings and had a diameter within 95–105% of the population average. Where destruction profiles are displayed, n is the number of oocytes from which the data have been gathered. Oocyte datasets were typically gathered from three independent experiments. For each independent experiment, both control and treatments groups were derived from the same pool of oocytes, collected from a minimum of two animals. Oocytes were selected at random for microinjection.

### Preparation of mRNA constructs for microinjection

H2B-mScarlet-hRec8(297-506)-mNeonGreen was a gift from Iain Cheeseman (Addgene plasmid # 174717 ; http://n2t.net/addgene:174717 ; RRID:Addgene_174717)^66^. Separase AA and the Scc1 and Rec8 separase sensors were constructed in pRN3 vectors as described previously^26^. Mouse *Sgo2la* cDNA (a gift from Mary Herbert) was amplified, and mutations were inserted, by primer overhang extension PCR. Following this, wild-type and mutant shugoshin-2 were inserted into a modified pRN3 vector designed to produce mRNA transcripts C-terminally coupled to mVenus^67^ using NEBuilder (New England Biolabs). Resultant plasmids were linearised and mRNA for microinjection was prepared using a T3 mMESSAGE mMACHINE kit (Ambion) according to the manufacturer’s instructions. mRNA was dissolved in nuclease-free water to the required micropipette concentration.

### Knockdown of gene expression by morpholino oligo

Morpholino antisense oligos designed to recognise the 5′-UTR of mouse securin (sequence: GATAAGAGTAGCCATTCTGGATTAC; MO; Gene Tools) or shugoshin-2 (sequence: CTGAAAACTAGTGGCGACCCCTCGC; MO; Gene Tools) were used to knock down gene expression. As per the manufacturer’s instructions, the oligo was stored at room temperature, heated for 5 minutes at 65°C prior to use, and injected at a micropipette concentration of 1 mM. Morpholinos stored in suspension for extended periods of time were autoclaved prior to use. All morpholinos were injected with sufficient time for protein turnover to result in an effective knock-down as quantified by western blot for securin^26^ and by immunofluorescence for shugoshin-2 (Supplementary Fig. 2).

### Microinjection

Oocyte microinjection of the MO and construct mRNAs was carried out on the heated stage of an inverted microscope fitted for epifluorescence (Olympus; IX71). In brief, micropipettes fabricated using a P-97 (Sutter Instruments) were inserted into cells using the negative capacitance overcompensation facility on an electrophysiological amplifier (World Precision Instruments). This procedure ensures a high rate of oocyte survival (>95%). The final volume of injection was estimated by the diameter of displaced ooplasm and was typically between 0.1 and 0.3% of total volume.

### Immunofluorescence

Cumulus-free oocytes synchronised in meiosis I by IMBX washout were attached to 8-well chamber slides using Cell-Tak (Corning) (all experimental groups were fixed and stained in the same well). Oocytes were washed with warm Phosphate Buffered Saline (PBS, Gibco) before being fixed for 30 minutes at room temperature in PBS containing 1.6% formaldehyde (Sigma Aldrich) and 0.1% Triton X-100 (Thermo Scientific). Following fixation, oocytes were further permeabilised in PBS containing 0.1% Triton X-100 at 4°C overnight before being washed into PBS containing 0.1% Tween (Sigma Aldrich) for storage of up to 2 days. Oocytes were blocked in 3% w/v Bovine Serum Albumin (BSA, Sigma Aldrich) PBS-Tween for 1-2 hours at room temperature, before incubating at 4°C overnight with primary antibodies. Shugoshin-2 knockdown was assayed with a polyclonal rabbit antibody from Biorbyt (orb499806, Lot: BS67182) used at 1:1000 in 3% BSA PBS-Tween. After 3x 5-minute washes in 3% BSA PBS-Tween, oocytes were incubated with donkey anti-rabbit Alexa Fluor Plus secondary antibodies at 1:1000 (Invitrogen). Samples were then washed twice with PBS-Tween and once with PBS before imaging.

### Microscopy

For all live-cell imaging, the far-red fluorescent DNA-intercalator SiR-DNA (Spirochrome) was added to the culture media 30 minutes prior to imaging at a final concentration of 125 nM. Media was maintained at 37°C under mineral oil. To generate destruction profiles of fluorescent constructs, Differential Interference Contrast (DIC) and fluorescence Z-sections were captured every 10 minutes through meiosis I using a Leica DMi8 inverted microscope with a K5 sCMOS camera (Leica Microsystems) and a 10x air objective lens. Fixed-cell immunofluorescence imaging was performed at room temperature using the same microscope and a 20x air objective lens.

Confocal images (all experiments using the separase biosensor) were captured using a Zeiss LSM 800 with a 40x oil immersion objective lens. Oocytes were imaged at 10-minute intervals through 20+ Z-sections over 12-15 hours from 10-20 minutes post nuclear envelope breakdown (NEBD). For separase sensor imaging, the laser power was adjusted so that the recorded intensity of the mCherry and eGFP channels were approximately equivalent. Transmitted light and fluorescent images were recorded in Zen Blue (Zeiss) and processed in the Fiji distribution of ImageJ^68^.

### Quantification and analysis

For real-time shugoshin-2 destruction profiling, images were recorded using LasX (Leica microsystems). Fluorescence intensity was subsequently measured in Fiji by taking a total VFP intensity reading from a defined region of interest around the oocyte. Data was plotted over time per oocyte aligned at PB1 extrusion. Fluorescence data values are arbitrary. Individual data sets were normalised prior to calculating the mean so that each data set was represented equally (taking the maximum fluorescence prior to anaphase as 100 arbitrary units). However, we also compared all raw traces, confirming that in each experiment, the timing and magnitude of construct destruction remained the same, regardless of the data handling method or variation in pre-destruction fluorescence. Comparisons typically include data from oocytes with comparable reporter expression levels. All mean destruction traces have associated data sets in individual oocytes.

For measuring shugoshin-2 knock-down in prometaphase I, a DNA clipping mask was generated using the SiR-DNA channel in Fiji. This mask was applied to the VFP channel to isolate DNA-localising signal and exclude non-specific primary antibody staining. The intensity of DNA-localising shugoshin-2 signal was then measured.

Cleavage profiles for separase biosensor experiments were produced in Fiji by again creating a clipping mask of the DNA using the SiR-DNA channel. The eGFP and mCherry signals within the area defined by the clipping mask were then measured and aligned at PB1 extrusion. Oocytes that did not undergo PB1 extrusion could not be aligned and were therefore excluded. eGFP/mCherry ratios were calculated in Excel. When plotting construct destruction traces or separase sensor ratios, for clarity, only the mean trace is shown with SEM error bars. For measuring chromosome alignment length, a maximum intensity projection of the SiR-DNA channel was produced in Fiji, and the distance between the furthest edges of the chromosomes was measured axial to the spindle.

### Multiple sequence alignments

Sequence conservation alignments were made by importing protein sequences from Uniprot and aligning in Jalview^69^, version 15.0. The full sequence alignment conservation annotation shown in Fig. 2a is between shugoshin-2 orthologs from *Homo sapiens*, *Mus musculus*, *Rattus norvegicus*, *Bos Taurus*, *Sus domesticus*, *Rhinolophus ferrumequinum*, and *Trachemys scripta elegans*. All figures were prepared in Adobe Illustrator CC, version 17.1.0.

## Data availability

All data supporting the findings of this study are available from the corresponding authors on request. Further information and requests for resources and reagents should be directed to and will be fulfilled by Suzanne Madgwick.

---

**Supplementary figure 1.**
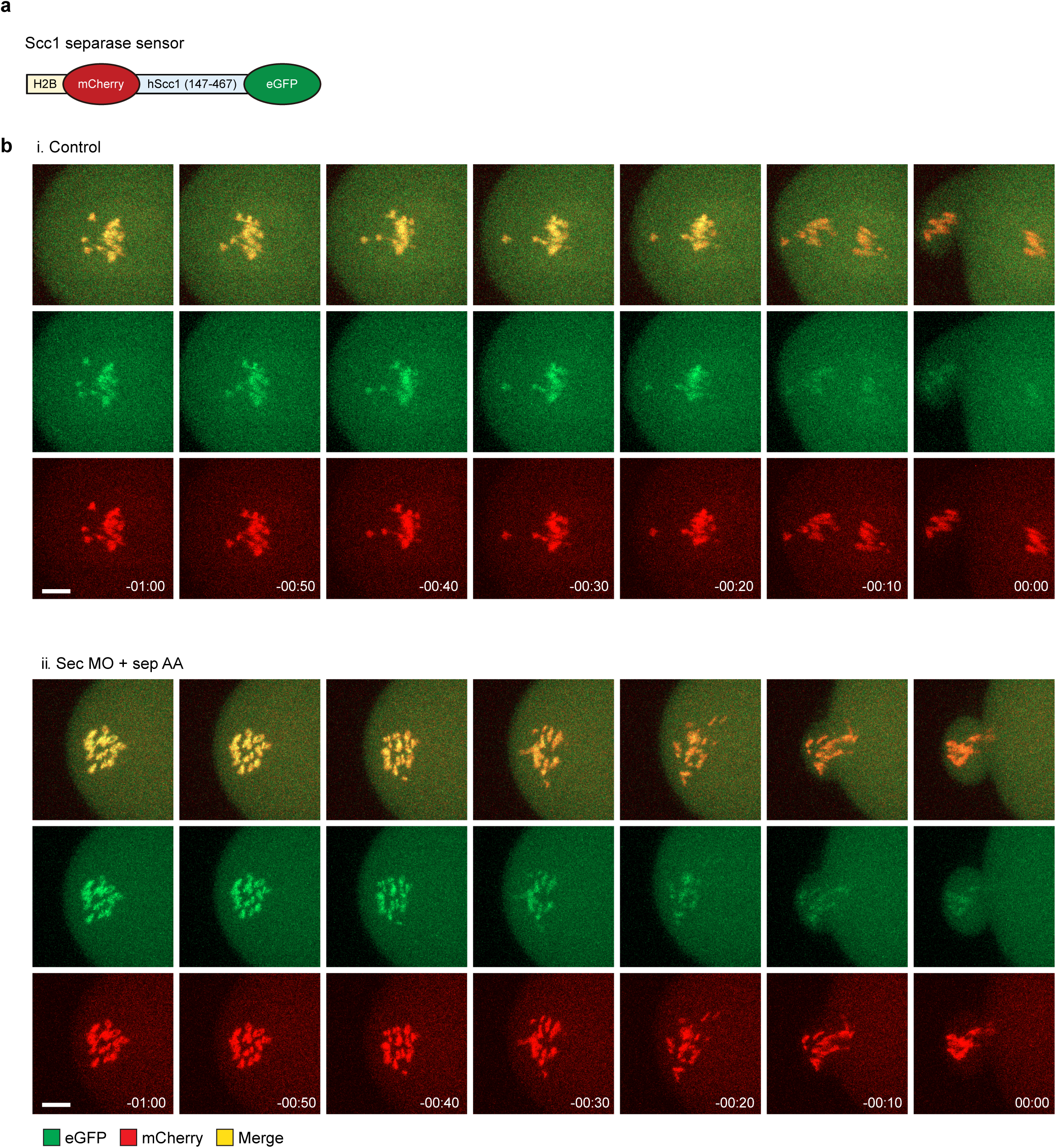
Separase sensor cleavage is detected earlier in oocytes where both securin and cyclin B1-CDK1 inhibitory pathways are restricted. (a) Schematic diagram showing the H2B-mCherry-hScc1-eGFP separase sensor. (b) Representaive images of (i) control and (ii) securin MO + separase AA oocytes at 10-minute intervals over 1 hour prior to PB1 extrusion. Panels show the mCherry signal which remains histone-bound, the eGFP signal which dissociates into the cytoplasm on sensor cleavage, and their overlay. Notably in (ii), securin MO + separase AA oocytes, the eGFP signal begins to decrease approximately 30 minutes earlier than in (i), control oocytes. In this example, the majority of chromosomes are extruded in the polar body.

**Supplementary figure 2.**
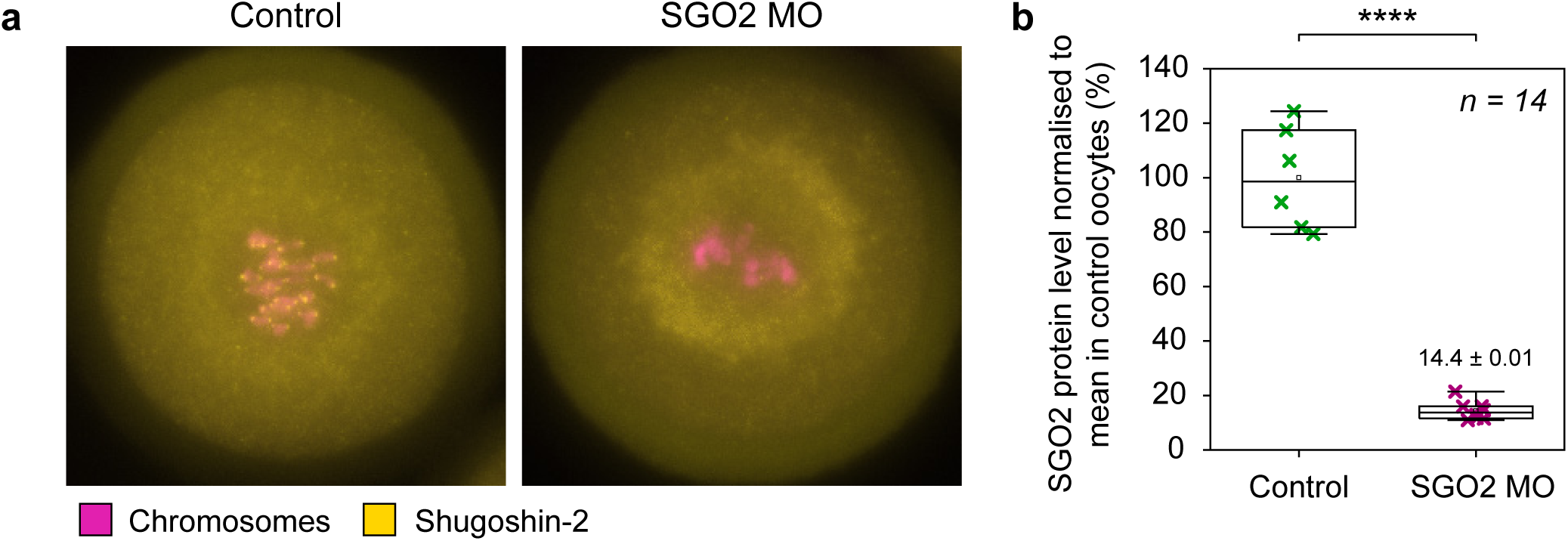
Quantification of shugoshin-2 knockdown by morpholino oligo. (a) Representative images of control and SGO2 MO oocytes collected at 3 hours post NEBD and immunostained for Shugoshin-2. (b) Quantification of shugoshin-2 protein levels in control (green crosses, n = 6) and SGO2 MO (magenta crosses, n = 8) oocytes normalised to the mean shugoshin-2 protein level in control oocytes. In panel (b): ****P<0.0001, ***P<0. 001, **P<0.01, *P<0.1, n.s. = non-significant. Significance was calculated by unpaired t-test.

